## Supplemental File 2 for "Personalizing chemotherapy drug selection using a novel transcriptomic chemogram"

| Gene | Frequency | Cis | Cyta | 5-FU | Gem | Irino | Lumi | Pac | Topo | Vinb | Vor |
|---|---|---|---|---|---|---|---|---|---|---|---|
| MMP10 | 6 | ✓ | ✓ | | | ✓ | | ✓ | ✓ | ✓ | |
| ADAT2 | 5 | ✓ | ✓ | ✓ | | | | ✓ | | ✓ | |
| NPM3 | 5 | ✓ | ✓ | ✓ | | | | ✓ | | ✓ | |
| C1QBP | 4 | ✓ | | ✓ | | | | ✓ | | ✓ | |
| RIOK1 | 4 | ✓ | | | | | | ✓ | | ✓ | ✓ |
| SLFN11 | 4 | ✓ | | | ✓ | ✓ | | | ✓ | | |
| KRT5 | 3 | ✓ | | | | | | ✓ | ✓ | | |
| LY6K | 3 | ✓ | | | | ✓ | | | ✓ | | |
| CLYBL | 3 | | ✓ | ✓ | | | | | | | ✓ |
| SLC27A5 | 3 | | ✓ | ✓ | | | | ✓ | | | |
| TAF4B | 3 | | ✓ | | | | | ✓ | | ✓ | |
| COQ3 | 3 | | | ✓ | | | | ✓ | | ✓ | |
| CSTA | 3 | | | ✓ | | | | ✓ | | ✓ | |
| FRAT2 | 3 | | | ✓ | | | | ✓ | | | ✓ |
| NRARP | 3 | | | ✓ | | | | ✓ | | | ✓ |
| SOX7 | 3 | | | | | ✓ | | ✓ | ✓ | | |
| WDFY2 | 3 | | | | | | ✓ | ✓ | ✓ | | |
| C15orf41 | 2 | ✓ | | | | | | | | ✓ | |
| CDCA7 | 2 | ✓ | | | | | | ✓ | | | |
| LRRC8C | 2 | ✓ | | | | | ✓ | | | | |
| PSAT1 | 2 | ✓ | ✓ | | | | | | | | |
| STOML2 | 2 | ✓ | | | | | | | | ✓ | |
| USP31 | 2 | ✓ | | | | | | ✓ | | | |
| ZNF750 | 2 | ✓ | | | | | | ✓ | | | |
| ACN9 | 2 | | ✓ | | | | | | | ✓ | |
| ASNS | 2 | | ✓ | | | | | | | ✓ | |
| CCNB1IP1 | 2 | | ✓ | | | | | | | ✓ | |
| F12 | 2 | | ✓ | ✓ | | | | | | | |
| FASTKD1 | 2 | | ✓ | ✓ | | | | | | | |
| MYC | 2 | | ✓ | | | | | ✓ | | | |
| POLR1D | 2 | | ✓ | ✓ | | | | | | | |
| SFXN4 | 2 | | ✓ | ✓ | | | | | | | |
| DSG3 | 2 | | | ✓ | | | | ✓ | | | |
| MYB | 2 | | | ✓ | | | | | | | ✓ |
| UQCRH | 2 | | | ✓ | | | | | | ✓ | |
| POLR3G | 2 | | | | ✓ | | ✓ | | | | |
| AIM2 | 2 | | | | | ✓ | | | ✓ | | |
| BNC1 | 2 | | | | | ✓ | ✓ | | | | |
| FOXL2 | 2 | | | | | ✓ | | | ✓ | | |
| ITPRIP | 2 | | | | | ✓ | ✓ | | | | |
| SERPINB4 | 2 | | | | | ✓ | | | ✓ | | |
| JARID2 | 2 | | | | | | ✓ | ✓ | | | |
| TAF5 | 2 | | | | | | | ✓ | | ✓ | |
| TMEM206 | 2 | | | | | | | ✓ | | ✓ | |
| ATP1B3 | 1 | ✓ | | | | | | | | | |
| CDC7 | 1 | ✓ | | | | | | | | | |
| FKBP14 | 1 | ✓ | | | | | | | | | |
| WDR3 | 1 | ✓ | | | | | | | | | |
| C12orf57 | 1 | | ✓ | | | | | | | | |
| DLEU1 | 1 | | ✓ | | | | | | | | |
| DPH5 | 1 | | ✓ | | | | | | | | |

| Gene | Frequency | Cis | Cyta | 5-FU | Gem | Irino | Lumi | Pac | Topo | Vinb | Vor |
|---|---|---|---|---|---|---|---|---|---|---|---|
| FAR1 | 1 | | ✓ | | | | | | | | |
| GNPNAT1 | 1 | | ✓ | | | | | | | | |
| MTHFD2 | 1 | | ✓ | | | | | | | | |
| NOB1 | 1 | | ✓ | | | | | | | | |
| SIGMAR1 | 1 | | ✓ | | | | | | | | |
| SNRPA1 | 1 | | ✓ | | | | | | | | |
| TUBE1 | 1 | | ✓ | | | | | | | | |
| ATP5D | 1 | | | ✓ | | | | | | | |
| CHCHD10 | 1 | | | ✓ | | | | | | | |
| FAM83F | 1 | | | ✓ | | | | | | | |
| GMDS | 1 | | | ✓ | | | | | | | |
| MRPL2 | 1 | | | ✓ | | | | | | | |
| MUC13 | 1 | | | ✓ | | | | | | | |
| PDSS1 | 1 | | | ✓ | | | | | | | |
| PIP5K1B | 1 | | | ✓ | | | | | | | |
| PPP1R1B | 1 | | | ✓ | | | | | | | |
| REG4 | 1 | | | ✓ | | | | | | | |
| RPL22L1 | 1 | | | ✓ | | | | | | | |
| SPINK4 | 1 | | | ✓ | | | | | | | |
| VSNL1 | 1 | | | ✓ | | | | | | | |
| ZNF511 | 1 | | | ✓ | | | | | | | |
| ARNTL2 | 1 | | | | ✓ | | | | | | |
| CRLF3 | 1 | | | | ✓ | | | | | | |
| CXCL1 | 1 | | | | ✓ | | | | | | |
| ELK3 | 1 | | | | ✓ | | | | | | |
| GLIPR1 | 1 | | | | ✓ | | | | | | |
| MLKL | 1 | | | | ✓ | | | | | | |
| PROCR | 1 | | | | ✓ | | | | | | |
| RELB | 1 | | | | ✓ | | | | | | |
| BCL2A1 | 1 | | | | | ✓ | | | | | |
| HMGA1 | 1 | | | | | ✓ | | | | | |
| PGM2 | 1 | | | | | ✓ | | | | | |
| SLC6A15 | 1 | | | | | ✓ | | | | | |
| TRAF3 | 1 | | | | | ✓ | | | | | |
| ADAMTS6 | 1 | | | | | | ✓ | | | | |
| ARHGAP22 | 1 | | | | | | ✓ | | | | |
| CDH13 | 1 | | | | | | ✓ | | | | |
| CSGALNACT2 | 1 | | | | | | ✓ | | | | |
| CTHRC1 | 1 | | | | | | ✓ | | | | |
| DSE | 1 | | | | | | ✓ | | | | |
| DYNC2H1 | 1 | | | | | | ✓ | | | | |
| DZIP1 | 1 | | | | | | ✓ | | | | |
| FAM101B | 1 | | | | | | ✓ | | | | |
| IL27RA | 1 | | | | | | ✓ | | | | |
| KRT14 | 1 | | | | | | ✓ | | | | |
| MFAP2 | 1 | | | | | | ✓ | | | | |
| PDLIM4 | 1 | | | | | | ✓ | | | | |
| POPDC3 | 1 | | | | | | ✓ | | | | |
| RFTN1 | 1 | | | | | | ✓ | | | | |
| SH3PXD2B | 1 | | | | | | ✓ | | | | |
| SLC31A2 | 1 | | | | | | ✓ | | | | |

| Gene | Frequency | Cis | Cyta | 5-FU | Gem | Irino | Lumi | Pac | Topo | Vinb | Vor |
|---|---|---|---|---|---|---|---|---|---|---|---|
| SLC4A7 | 1 | | | | | | ✓ | | | | |
| SPHK1 | 1 | | | | | | ✓ | | | | |
| TM4SF19 | 1 | | | | | | ✓ | | | | |
| TWIST1 | 1 | | | | | | ✓ | | | | |
| VEGFC | 1 | | | | | | ✓ | | | | |
| IL1B | 1 | | | | | | | | ✓ | | |
| RAB38 | 1 | | | | | | | | ✓ | | |
| C6orf170 | 1 | | | | | | | | | ✓ | |
| ECSIT | 1 | | | | | | | | | ✓ | |
| FGF11 | 1 | | | | | | | | | ✓ | |
| FSD1 | 1 | | | | | | | | | ✓ | |
| LRRC49 | 1 | | | | | | | | | ✓ | |
| NOC3L | 1 | | | | | | | | | ✓ | |
| RPF2 | 1 | | | | | | | | | ✓ | |
| SEH1L | 1 | | | | | | | | | ✓ | |
| TBPL1 | 1 | | | | | | | | | ✓ | |
| ARID3B | 1 | | | | | | | | | | ✓ |
| C9orf152 | 1 | | | | | | | | | | ✓ |
| CECR5 | 1 | | | | | | | | | | ✓ |
| DCXR | 1 | | | | | | | | | | ✓ |
| DQX1 | 1 | | | | | | | | | | ✓ |
| EFNA3 | 1 | | | | | | | | | | ✓ |
| FHIT | 1 | | | | | | | | | | ✓ |
| FKBP4 | 1 | | | | | | | | | | ✓ |
| IL17RB | 1 | | | | | | | | | | ✓ |
| JHDM1D | 1 | | | | | | | | | | ✓ |
| NOC2L | 1 | | | | | | | | | | ✓ |
| PDCD2L | 1 | | | | | | | | | | ✓ |
| PEX7 | 1 | | | | | | | | | | ✓ |
| PHGR1 | 1 | | | | | | | | | | ✓ |
| SDHAF1 | 1 | | | | | | | | | | ✓ |
| TIMM8B | 1 | | | | | | | | | | ✓ |
| TMEM168 | 1 | | | | | | | | | | ✓ |
| TMEM183A | 1 | | | | | | | | | | ✓ |
| TOX3 | 1 | | | | | | | | | | ✓ |
| TRAP1 | 1 | | | | | | | | | | ✓ |
| TTC39A | 1 | | | | | | | | | | ✓ |
| AMTN | 1 | | | | | | | ✓ | | | |
| ARTN | 1 | | | | | | | ✓ | | | |
| C1orf74 | 1 | | | | | | | ✓ | | | |
| CYB5R4 | 1 | | | | | | | ✓ | | | |
| FAT2 | 1 | | | | | | | ✓ | | | |
| GBP6 | 1 | | | | | | | ✓ | | | |
| GNA15 | 1 | | | | | | | ✓ | | | |
| GPC2 | 1 | | | | | | | ✓ | | | |
| HOXD10 | 1 | | | | | | | ✓ | | | |
| KRT6A | 1 | | | | | | | ✓ | | | |
| KRT6B | 1 | | | | | | | ✓ | | | |
| MARK1 | 1 | | | | | | | ✓ | | | |
| MMP13 | 1 | | | | | | | ✓ | | | |
| PKP1 | 1 | | | | | | | ✓ | | | |

| Gene | Frequency | Cis | Cyta | 5-FU | Gem | Irino | Lumi | Pac | Topo | Vinb | Vor |
|---|---|---|---|---|---|---|---|---|---|---|---|
| PREP | 1 | | | | | | | ✓ | | | |
| PVRL1 | 1 | | | | | | | ✓ | | | |
| REL | 1 | | | | | | | ✓ | | | |
| S100A7 | 1 | | | | | | | ✓ | | | |
| SH3BP1 | 1 | | | | | | | ✓ | | | |
| TP63 | 1 | | | | | | | ✓ | | | |
| TRERF1 | 1 | | | | | | | ✓ | | | |
